# Gut microbiota of dogs with cancer receiving anti-EGFR/HER2 immunization reveals potential biomarkers of patient survival

**DOI:** 10.1101/2025.07.13.663784

**Authors:** Richard R Rodrigues, Vini Karumuru, Stephanie Nuss, Marina Elliott, Isaiah Shriver, Chih-Min Chao, Hester A Doyle, Chelsea Tripp, Mark J Mamula, Amiran Dzutsev, Andrey Morgun, Natalia Shulzhenko

## Abstract

**Background:** Canine cancer remains a leading cause of death in dogs, yet advances in veterinary oncology lag behind human medicine, particularly in immunotherapy. While immune checkpoint inhibitors are just entering clinical trials in dogs, other immunotherapies, such as anti-EGFR/HER2 vaccines, have shown promise. In parallel, mounting evidence in human oncology links gut microbiota composition to immunotherapy response. However, this relationship remains unexplored in canine patients. In this pilot study, we analyzed the gut microbiome of dogs enrolled in a clinical trial of anti-EGFR/HER2 immunotherapy to identify microbial biomarkers associated with survival outcomes.

**Methods:** Rectal swab samples of 51 dogs were collected at the time of first vaccine administration (baseline microbiota) and underwent 16S rRNA gene sequencing according to standard protocols.

**Results:** Microbiome composition showed no significant differences by cancer type, sex, or breed, suggesting no inherent microbiome bias in the cohort. However, Cox regression analysis revealed 11 bacterial taxa whose abundances were significantly associated with overall survival (FDR < 0.1), independently of cancer type. Seven taxa were linked to increased mortality risk, while four were associated with prolonged survival. These associations remained significant after adjusting for confounders such as hemangiosarcoma diagnosis and advanced age.

**Conclusions:** To our knowledge, this is the first study to identify gut microbial signatures associated with survival in dogs undergoing cancer immunotherapy. These findings suggest that specific bacterial taxa may serve as prognostic biomarkers for immunotherapy outcomes in canine cancer, laying the groundwork for microbiota-targeted strategies to improve therapeutic efficacy in veterinary oncology.

## BACKGROUND

Cancer remains the leading cause of mortality in dogs, yet veterinary oncology lags behind advancements achieved in human cancer therapies. While checkpoint inhibitors—revolutionary treatments in human oncology—have only recently entered canine clinical trials, other immunotherapies for dogs have advanced over the past decade [1]. For example, a novel therapy designed to induce EGFR/HER2-specific immunity demonstrated promising results in stimulating T and B cell anti-tumor activity and reducing metastatic tumor burden in canine patients [2]. These developments underscore the growing potential of immunotherapy in veterinary medicine, though challenges in translating human innovations persist.

Parallel to immunotherapy progress, decades of immunological research have established the critical role of the gut microbiota—the community of commensal microbes inhabiting the gastrointestinal tract—in modulating both local (intestinal) and systemic immune responses [3]. This connection has positioned gut microbiota as a key focus in oncological research. Consequently, microbiota-modifying therapies, such as fecal microbiota transplants (FMT), are now being explored in clinical trials as adjuvants to enhance immunotherapy efficacy [4]. Notably, studies in humans have linked specific gut microbiome signatures to clinical outcomes in cancer patients receiving immunotherapy highlighting their potential as diagnostic or prognostic biomarkers [5, 6].

Building on this foundation, our study is the first to investigate the gut microbiota of canine cancer patients undergoing anti-cancer immunotherapy, namely anti-EGFR immunization. While the baseline gut microbiome composition did not differ significantly across cancer types, we identified eleven bacterial taxa whose abundances correlated with patient survival, independent of cancer type. These findings provide preliminary evidence that specific gut microbes may serve as prognostic markers for immunotherapy response in dogs. This discovery aligns with emerging trends in human oncology and suggests that microbiota-targeted strategies could enhance therapeutic outcomes in veterinary cancer care.

## METHODS

### Animals

Canine cancer patients were recruited at Bridge Animal Referral Center (Edmonds, WA) as part of an ongoing clinical trial of anti-EGFR/HER2 immunization of dogs with cancer [2]. Rectal swabs were collected with the approval and informed consent of the canine patient owners on the day of the first vaccine administration (baseline microbiome). Protocols were consistent with accepted guidelines of the NIH for the care and use of animals as well as approved by the Yale University Institutional Animal Care and Use Committee.

All patients received standard of care recommended by a veterinary oncologist. Two injections of EGFR/HER2 peptide vaccine administered 3 weeks apart as previously described [2].

A total of 51 dogs were included in this pilot study with 28 (54.9%) females, 34 (66.7%) pure breed dogs, mean age at diagnosis 9.5 years (SD 2.6 years). The most frequent diagnoses in this cohort were osteosarcoma (OSA, n=22), followed by hemangiosarcoma (HAS, n=8). Several other types of cancers with fewer than five dogs were combined as other cancers (OthCA, n=21). Detailed demographic information for each dog can be found in Supplementary Table 1.

### Samples

Rectal swabs for microbiome analysis were collected using Zymo Research DNA/RNA Shield Safe Collect Swab Collection kit (1ml, Cat. #R1160-E) and stored at +4oC until shipped to the lab for processing.

DNA was extracted using ZymoBIOMICs DNA/RNA Miniprep Kit (Cat. # R2002) according to the manufacturer instructions with the beat beating done in BeadRuptor (OMNI).

Extracted DNA underwent a two-step PCR protocol to amplify and barcode the 16S rRNA gene V4 region (515F-806R) at the National Cancer Institute Microbiome and Genetics Core Facility using their standard workflow [7]. Samples were sequenced using 250-bp paired read run on the Illumina MiSeq platform.

### Data Analysis

The analysis of the 16S rRNA gene sequencing data was done as previously described [7]. Paired-end fastq reads were analyzed using DADA2 [8] and QIIME2 2019.4 [9]. The 51 samples contained an average of 32,262 reads per sample, and a total of 899 ASVs were detected.

The downstream analysis of the microbiome abundance data was done using the JAMS package version 1.9.8 with R/4.4.1, available at https://github.com/johnmcculloch/JAMS_BW. For comparisons between samples, ordination plots based on of last known taxon (LKT) abundances [5] were made with the t-UMAP algorithm using the uwot package in R (https://github.com/jlmelville/uwot) and the ggplot2 library. PERMANOVA values were obtained using the adonis function of the vegan package, with default (999) permutations and pairwise distances calculated using Bray–Curtis distance. Functional (EC, KEGG and MetaCyC) information in the metagenomic samples were predicted using PICRUSt2 (v 2.6.0) [10] by running the full default pipeline via the picrust2_pipeline.py command along with the PICRUSt2-SC database [11]. The analysis of the overall survival data (time and outcome) and covariates was done as previously described [5]. For biomarkers with continuous values, cutoff points were computed with the ‘cutp’ function (survMisc v0.5.5 package) and samples were categorized into high and low groups based on the biomarker values.

Kaplan–Meier plots were generated using a combination of survival v3.2-11, ggplot2 v3.3.5 and plotly packages. Finally, univariate and multivariate Cox regression analyses for categories were performed using the coxph function from survival package, and hazard ratios (HRs) and P values were calculated.

Data were submitted to NCBI SRA database under BioProject ID PRJNA1276127.

## RESULTS

In the initial phase of this study, we sought to broadly characterize gut microbial communities in relation to key clinical factors including cancer type, sex, and breed. Neither alpha diversity (within-sample richness and evenness) nor beta-diversity (between-sample compositional differences) metrics exhibited significant differences with respect to any of these factors (Figure 1, Suppl Table 2).

**Figure 1.**
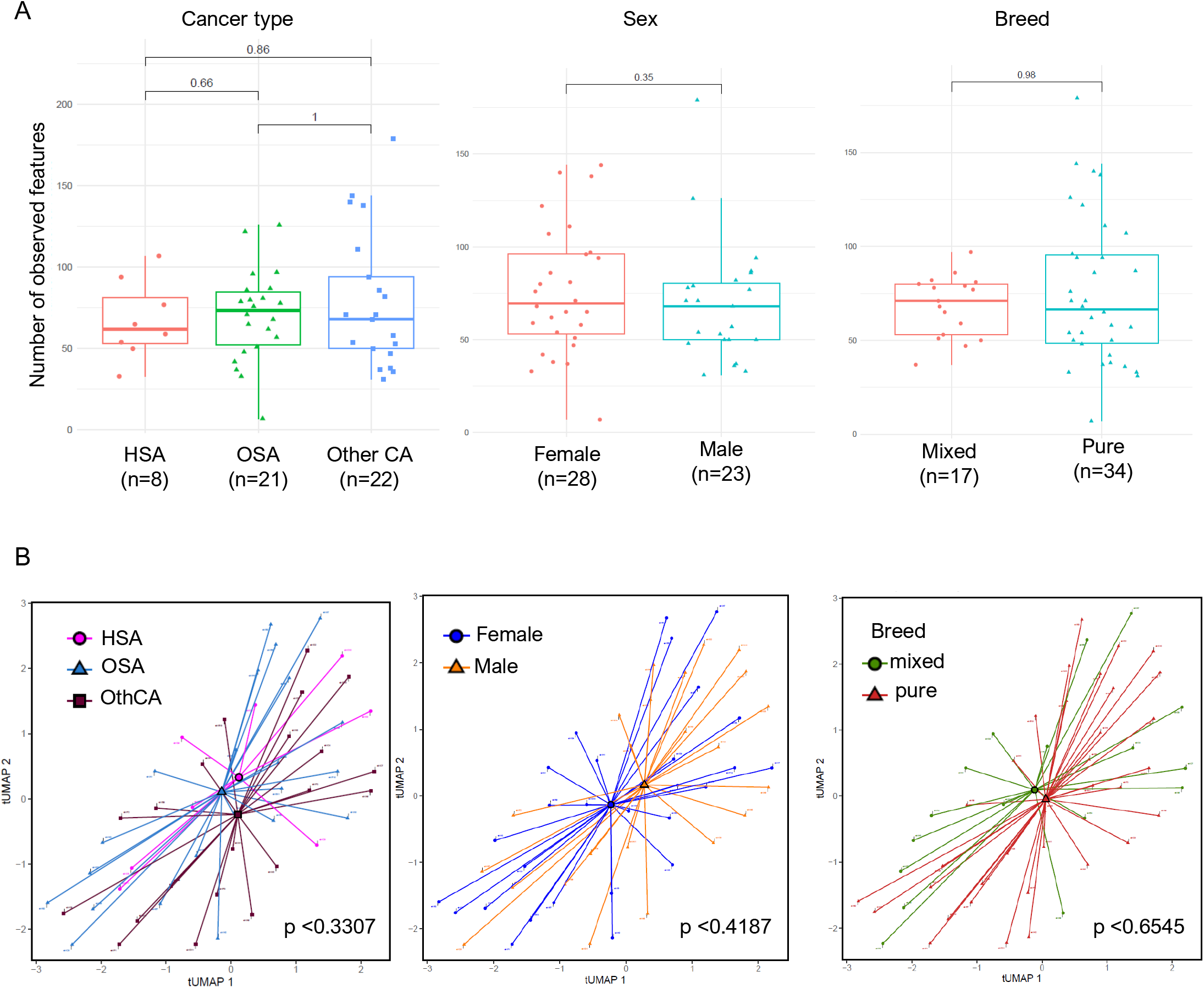
Microbiome diversity metrics in 51 dogs with cancer in this study. **A**. Microbiome alpha diversity compared between dogs with hemangiosarcoma (HSA), osteosarcoma (OSA), other cancers (OthCA), dogs of different sexes and breeds showing no difference between the groups. **B**. Ordination t-UMAP plots of Bray-Curtis dissimilarity indexes showing no compositional differences between patients of different cancer types, sexes or breeds. All comparisons are based on last known taxon (LKT) abundances, two-tailed P values calculated using PERMANOVA.

Consistent with the absence of differences in beta-diversity, there were no statistically significant differences in between the abundance of specific microbial taxa or their predicted functional categories associated with the aforementioned variables (data not shown). These results suggest that, within the limitations of our study (sample size and cohort characteristics), the gut microbiome composition may lack sufficient specificity for distinguishing cancer types in dogs. However, it also indicated that there was no inherent bias in the study population.

Next, we focused on the study’s primary objective: identifying microbial taxa with the potential of being biomarkers of canine cancer survival. To address this, we first evaluated which clinical variables were associated with the survival of canine patients. By assessing sex, breed, cancer type, age, and weight, we found that dogs with HSA as well as those older than 11.9 years at diagnosis had lower survival rates compared to other patients, whereas breed did not make a difference for survival (Figure 2). These findings align with prior reports of poor survival in older canines and those with HSA, reinforcing the consistency of our cohort and underscoring key confounding factors that could influence analyses of microbiota composition-survival relationships.

**Figure 2.**
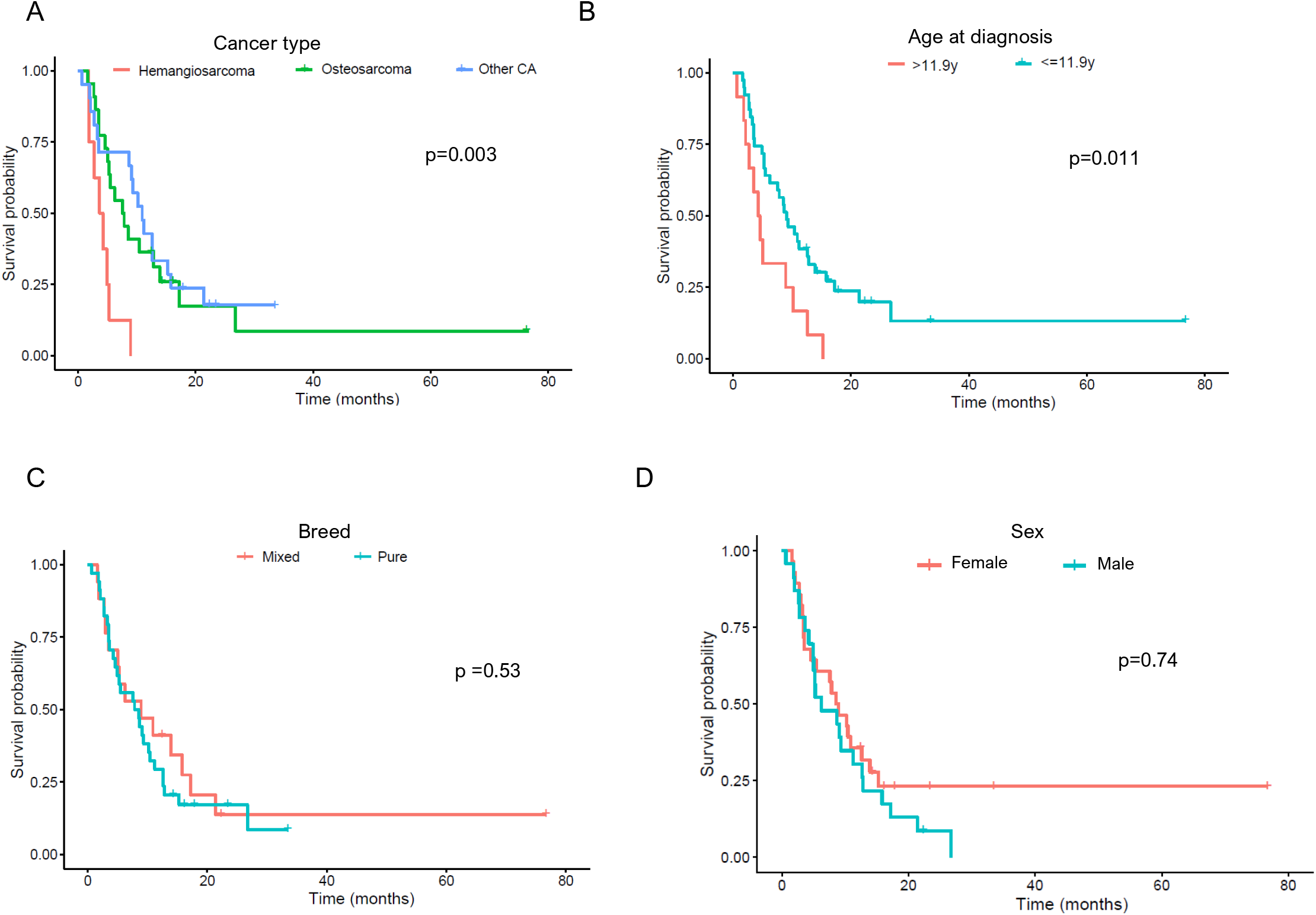
Kaplan-Meier curves of survival probability of dogs with cancer stratified by cancer type (A), age (B), breed (C) and sex (D). Numbers of patients at risk at each time point are shown. Score (log rank) test two-tailed P value from Cox proportional hazards regression analysis are shown.

Given that cancer type emerged as the most significant factor in our analysis, with HSA demonstrating the poorest survival outcomes and smallest sample size, we evaluated microbiome-survival relationships in the other groups: OSA (n=22) and OthCA (n=21). Our goal was to determine whether microbial associations with survival were consistent across cancer types. Using Cox regression under a relaxed false discovery rate (FDR) threshold of <0.25, we identified 57 taxa associated with survival in OSA and 46 taxa in OthCA (Figure 3).

**Figure 3.**
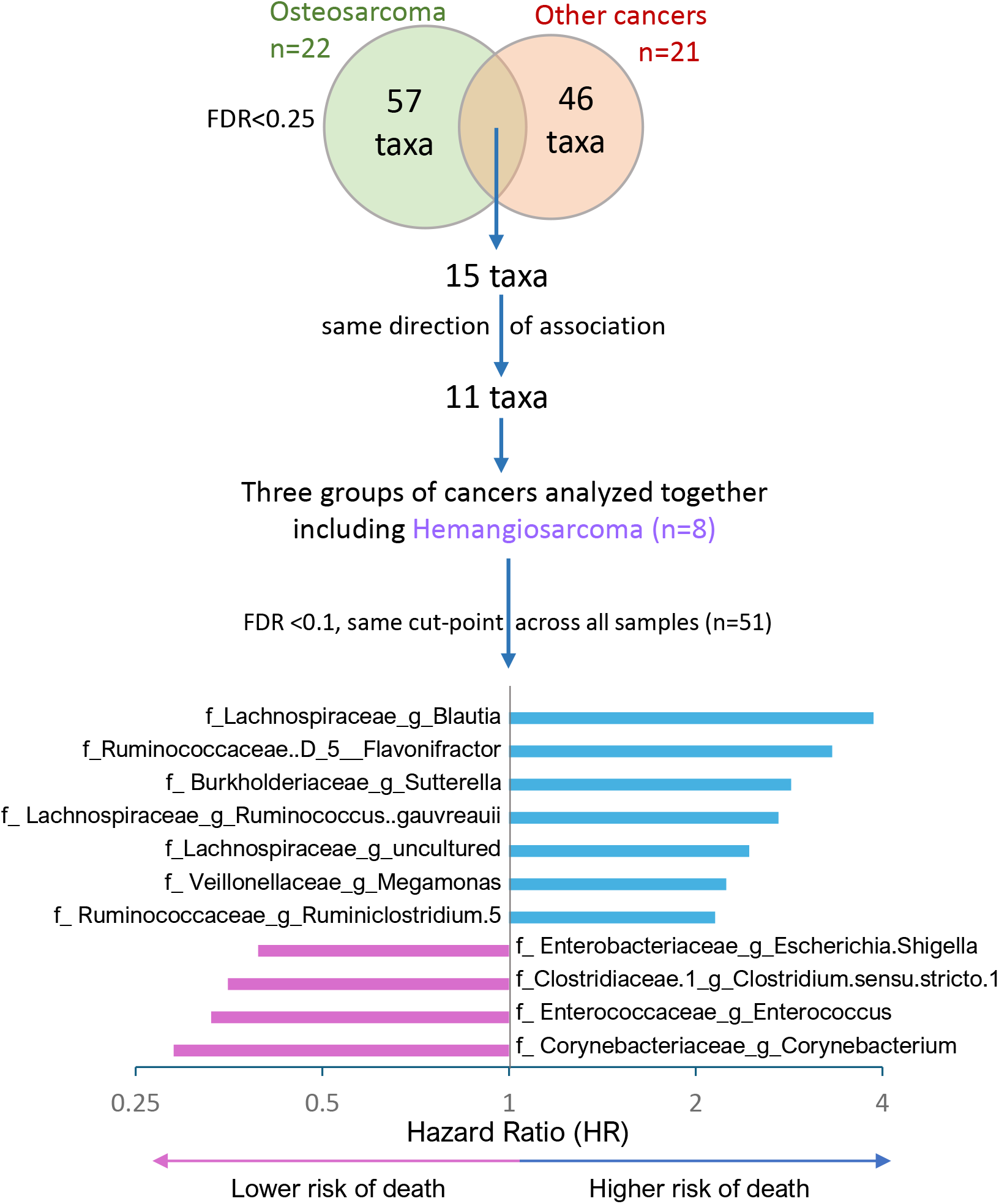
Workflow of analysis for identification of microbial taxa associated with canine patient survival. Cox regression analysis was used to identify taxa associated with survival in the indicated types of cancers. FDR, false discovery rate.

Among the 15 taxa overlapping between these two groups, 11 microbes (73%) exhibited concordant directions of association (i.e., hazard ratios), suggesting shared survival-related microbial signatures across cancer types. Notably, while these microbes were common to both groups, the optimal abundance cut-points (low vs. high) differed between OSA and OthCA patients.

We then performed Cox regression on the combined set of all 51 dogs to include HSA. All 11 previously identified taxa remained significantly associated with survival under stricter statistical criteria (FDR<0.1), with 7 taxa linked to the increased risk of death (Hazard Ratio (HR)>1) and 4 taxa associated with longer dog survival (HR<1; Figure 3). Importantly, the same abundance thresholds (low/high) were applied uniformly across the entire cohort of dogs to enable direct comparison across all samples represented by the survival curves based on the abundances of taxa positively (Figure 4) and negatively (Figure 5) associated with dog survival.

**Figure 4.**
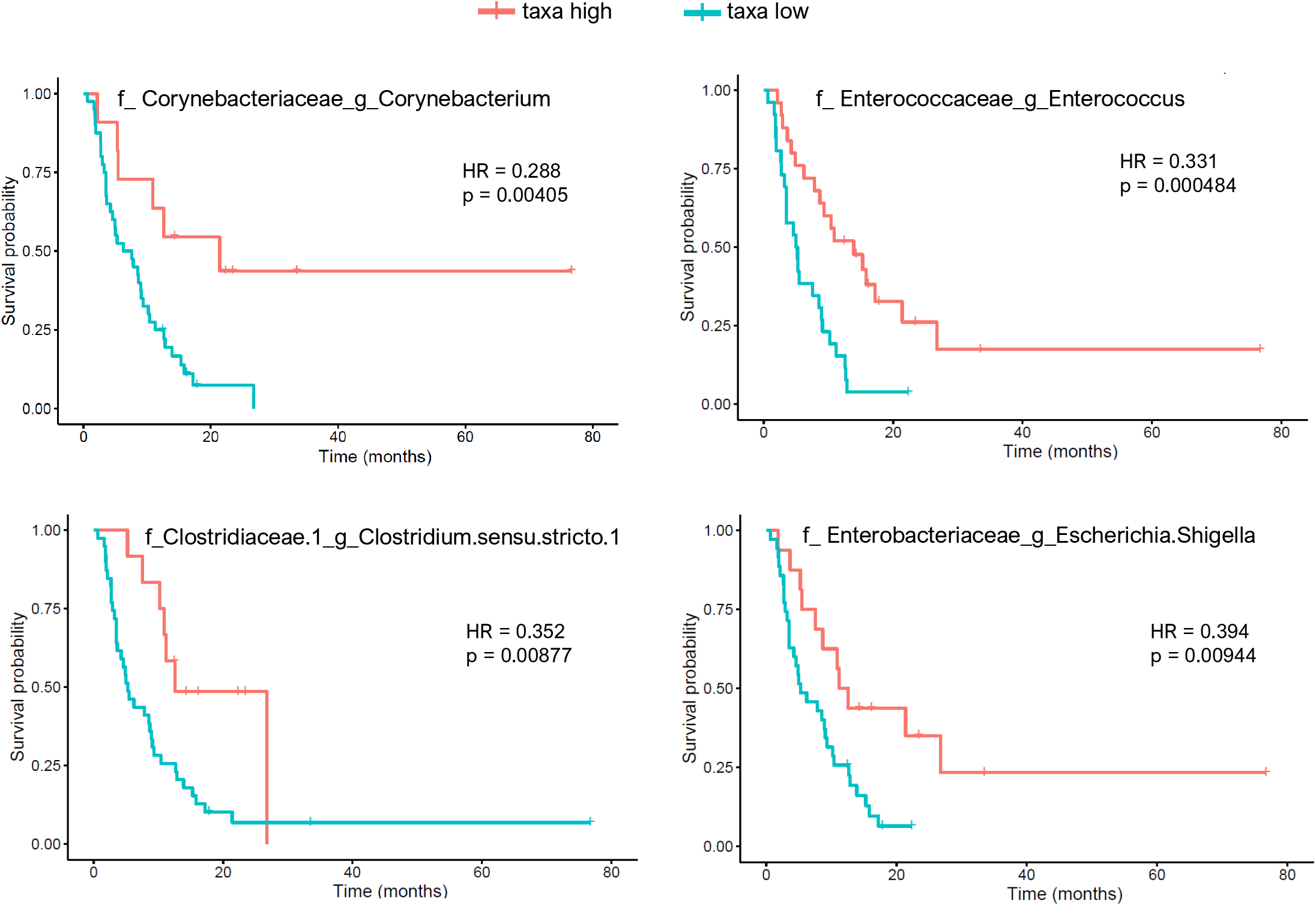
Kaplan-Meier curves of survival probability of dogs (n=51) based on relative abundances of taxa **positively** associated with survival (Hazard Ratio (HR) of death <1). Taxa identified by the family (f) and genus (g) ranks. Samples were classified as ‘taxa high’ or ‘taxa low’ based on whether the relative abundance was above or below the cut-point threshold. Score (log rank) test two-tailed p values from Cox proportional hazards regression analysis are shown. Full taxonomic identification and cut-points for each taxon is in Supplementary Table 3.

**Figure 5.**
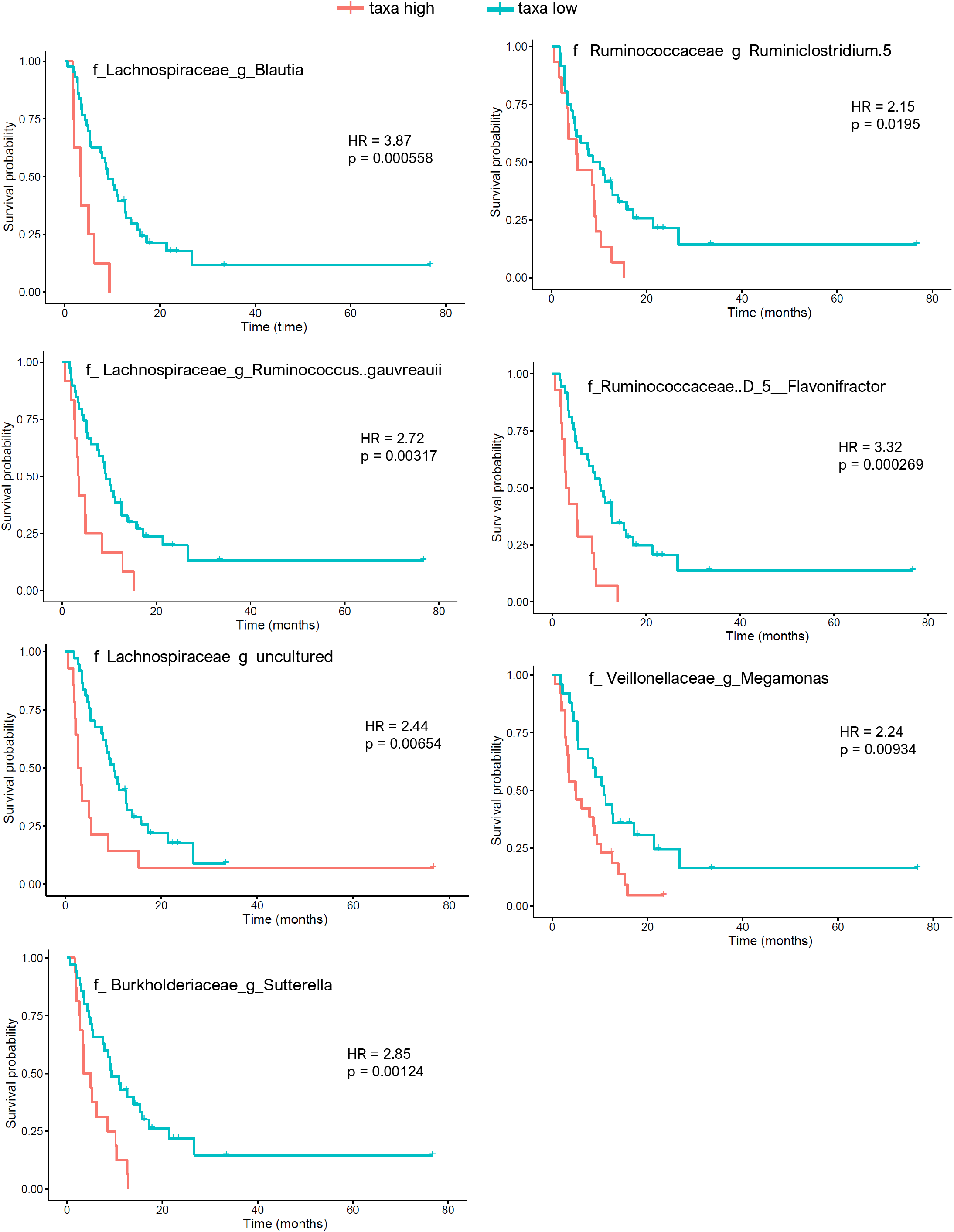
Kaplan-Meier curves of survival probability of dogs (n=51) based on relative abundances of taxa **negatively** associated with survival (Hazard Ratio (HR) of death >1). Taxa identified by the family (f) and genus (g) ranks. Samples were classified as ‘taxa high’ or ‘taxa low’ based on whether the relative abundance was above or below the cut-point threshold. Score (log rank) test two-tailed p values from Cox proportional hazards regression analysis are shown. Full taxonomic identification and cut-points for each taxon is in Supplementary Table 3.

Finally, given the markedly poor survival observed in HSA patients and dogs older than 11.9 years, we explicitly tested whether these factors confounded the observed microbial associations. After adjustment, all 11 microbes retained significant associations with survival (FDR<0.1), confirming their robustness to these covariates (Suppl. Table 3).

## DISCUSSION

Following the success in identifying microbiome signatures predictive of survival in human cancer patients, this pilot study aimed to examine this question for canine cancers. Accordingly, we identified eleven gut microbial taxa robustly associated with survival in dogs undergoing anti-EGFR immunotherapy.

Several studies have investigated gut microbiota in dogs with cancer [12-14]. However, most of this work inquired about differences in microbiome between healthy and diseased dogs with different cancer types. Only one study attempted to analyze microbiome in relation to canine patient survival in 23 dogs with diverse cancers but the analysis was very limited [15].

Thus, to the best of our knowledge, this study is the first to link specific gut microbiota taxa to outcomes in canine cancer during immunotherapy. It is worth noting that these findings were independent of cancer type, age, or poor-prognosis factors such as hemangiosarcoma. This suggests that those microbes may serve as biomarkers of response to immunotherapy, rather than indicators of overall cancer survival irrespective of treatment type. Moreover, while we found promising associations, the results must be interpreted cautiously.

In contrast to veterinary field, human oncology has a wealth of research exploring microbiome signatures linked to survival and immunotherapy responses [5, 16, 17]. However, directly comparing our results with human literature remains challenging for a couple of reasons: First, the canine cohort we studied received a unique anti-EGFR vaccine treatment, which differs significantly from typical human cancer immunotherapies. Second, standard 16S rRNA gene sequencing often cannot identify bacteria at the species or strain level. These specific classifications can be crucial because they’re more likely to share the same functional characteristics.

Therefore, it’s not surprising that some bacterial genera we identified as being associated with survival in our canine study, such as Enterococcus and Ruminiclostridium, showed divergent findings with some human studies [17]. However, other genera like Flavonifractor, Lachnospiraceae, and Shigella have been reported with similar outcomes concerning the microbiome’s role in human cancer immunotherapy [17, 18].

Despite these intriguing parallels and divergences, it’s crucial to acknowledge the inherent limitations of this study, which can guide our interpretations and future research directions.

First, the study design does not clearly discriminate whether the associated taxa are biomarkers of cancer survival in general or specifically tied to the immunotherapy used here. Second, the relatively small sample size prevented us from identifying biomarkers specific to each cancer type. While this is a promising initial study, establishing reliable biomarkers and/or predictive tools will require not only larger sample sizes to narrow confidence intervals but also validation across multiple independent patient cohorts, which our collaborative group is currently pursuing.

Another direction for future studies is to explore the potential biological role of these microbes. Indeed, a substantial body of literature demonstrates that microbes can both enhance and diminish the effects of immunotherapy in animal models (such as mice) and human patients. Therefore, in addition to repeating this study with larger sample size and in independent cohorts, it is necessary to investigate the potential mechanistic (i.e., causal) role of gut microbiota in canine patient survival. To start, methodologies for causal discovery from observational data, like Mendelian Randomization [19], Transkingdom Network Analysis [20], and Mediation Analysis [21], could be employed to pinpoint microbes that are not just associated with better survival but whose supplementation could alter the disease course in canine patients. Next step would be implementing interventional causality approaches, such as randomized clinical trials involving fecal microbiota transplants and/or anti-cancer probiotics.

## CONCLUSIONS

Overall, this research establishes eleven gut microbial taxa as promising indicators of survival outcomes in dogs treated with anti-EGFR immunotherapy, even after accounting for diverse confounders, while finding no significant microbiome diversity correlations with key clinical factors. These results highlight the microbes’ role as potential immunotherapy-specific biomarkers, extending insights from human and veterinary studies. However, a relatively small cohort size limits the generalizability of these results, reflecting the preliminary status of our findings. Future efforts should prioritize larger, independent cohorts for developing reliable biomarkers. By addressing these gaps, we can advance microbial-based diagnostics and personalized medicine, ultimately enhancing survival prospects for canine cancer patients that can potentially benefit human oncology.

## List of abbreviations

HSA: Hemangiosarcoma
OSA: Osteosarcoma
OthCA: Other cancer
HR: Hazard Ratio
FDR: false discovery rate

## Availability of data and materials

Data are available at the NCBI SRA database under BioProject ID PRJNA1276127.

Competing interests None

## Funding

Canine Cancer Alliance, NIH NCI Intramural Program

## Authors’ contributions

RR, AD, AM, NS contributed to the conception and design of the work; VK, SN, ME, IS, HD, CT, MM contributed to data acquisition; RR, VK, CC, AD, AM, NS - data analysis and interpretation; RR, NS, AM - drafted the manuscript. All authors have approved the submitted version.

## Acknowledgements

The authors would like to thanks participants of the study, Bridge Animal Referral Center personnel for sample collection and the Canine Cancer Alliance for support.

## Notes

https://www.ncbi.nlm.nih.gov/bioproject/?term=PRJNA1276127

### Competing Interest Statement

The authors have declared no competing interest.

